# Syringeal vocal folds do not have a voice in zebra finch vocal development

**DOI:** 10.1101/2020.09.14.295931

**Authors:** Alyssa Maxwell, Iris Adam, Pernille S. Larsen, Peter G. Sørensen, Coen P.H. Elemans

## Abstract

Vocal behaviour can be dramatically changed by both neural circuit development and postnatal maturation of the body. During song learning in songbirds, both the song system and syringeal muscles are functionally changing, but it is unknown if maturation of sound generators within the syrinx contributes to vocal development. Here we densely sample the respiratory pressure control space of the zebra finch syrinx *in vitro*. We show that the syrinx produces sound very efficiently and that key acoustic parameters, minimal fundamental frequency, entropy and source level, do not change over development in both sexes. Thus, our data suggests that the observed acoustic changes in vocal development must be attributed to changes in the motor control pathway, from song system circuitry to muscle force, and not by material property changes in the avian analog of the vocal folds. We propose that in songbirds, muscle use and training driven by the sexually dimorphic song system are the crucial drivers that lead to sexual dimorphism of the syringeal skeleton and musculature. The size and properties of the instrument are thus not changing, while its player is.

## Introduction

Many vertebrates change their vocal output over postnatal development^1–4^. Especially in species capable of vocal imitation learning, progressive changes in vocal output are attributed to changes in the underlying brain circuitry. However, the maturation of the vocal organ and tract have profound effect on voice during puberty in humans^5^, and can explain transitions from juvenile to adult calls in marmosets^6^. Thus, the postnatal development of the body can drive changes in vocal behavior, and therefore needs to be included when studying vocal development.

The zebra finch is a widespread animal model to study the mechanisms underlying vocal imitation learning and production^7,8^. In male zebra finches, sound production changes dramatically over the 100-day period of from hatching to adulthood^2,9,10^, developing from variable subsong to highly stereotyped crystallized song. These changes, including the ones in acoustic structure, are typically attributed to changes in neural circuitry, even though the vocal organ, the syrinx, is also undergoing significant anatomical and functional changes during this time^11–14^. Coinciding with the transition from the sensory to the sensorimotor phase of learning around 20 DPH, syringeal muscles in male zebra finches increase in mass and cross-sectional area^11^. Additionally, the contractile properties of male syringeal muscles increase in speed^12^, which affects the transformation from neural commands to muscle force^14^. Thus, over development functional changes occur in the syrinx that change the transformation from neural signals into force which thereby change motor control of the vocal organ. However, it remains unknown if the organ itself also exhibits changes that will affect the complex transformation from muscle forces to sound.

Analogous to mammals, birds produce sound when expiratory airflow from the bronchi induces self-sustained vibrations of vocal fold-like structures within the syrinx^15–18^ (Fig. 1a) following the myoelastic-aerodynamic (MEAD) theory for voice production^19^. Songbirds have two bilateral sets of independently controlled paired folds or labia, one in each bronchus or hemi-syrinx, with the lateral labium on the lateral side, and medial vibratory mass (MVM) on the medial side. The MVM is a tissue continuum in which the thicker part is called the medial labium (ML)^20–22^ (Fig. 1b). These labia consist of multiple layers of tissue in most species investigated^22,23^. According to MEAD, acoustic features, such as fundamental frequency (*f*_o_), or boundary conditions for vibration, such as the phonation threshold pressure, are largely set by the positioning of and driving pressures^19^ on the vocal folds, as well as their resonance properties and shape. Vocal fold resonance properties in turn are determined by vocal fold length and ultrastructure, such as collagen and elastin fiber composition, the occurrence of tissue layering and layer orientation^24^. During vocal development in humans^25–27^, and marmosets^6^, changes in size, (ultra)structure and mechanical properties of the labia have been linked to dramatic changes in vocal output. However, it is unknown whether material properties or morphology of songbird labial change over vocal development and if so whether these changes contribute to the changing vocal output.

**Figure 1.**
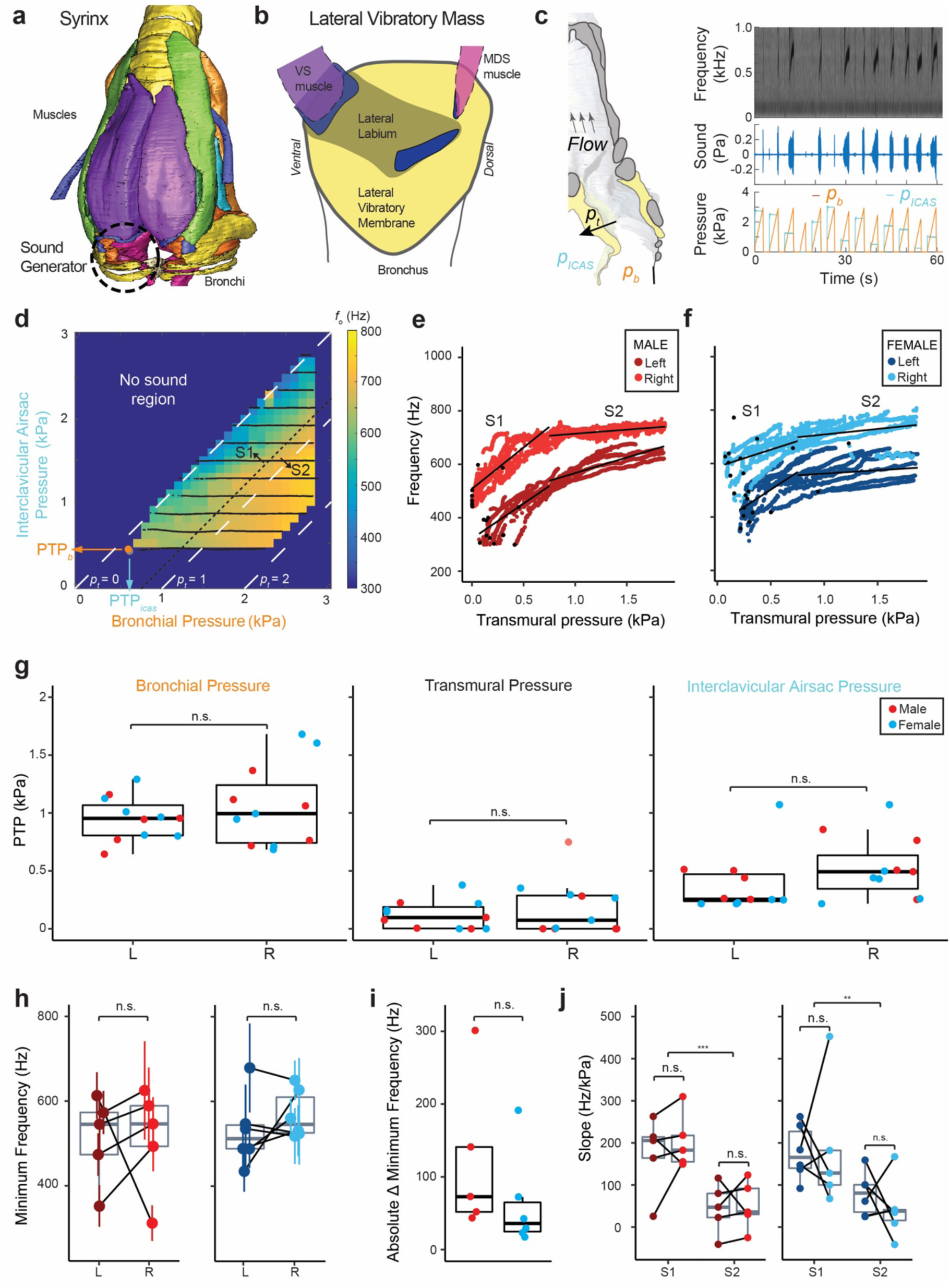
The adult zebra finch syrinx produces sound in a well-defined pressure control space with higher frequencies on the right. **(a)** Ventral view of the male syrinx. One of the two sound generators are indicated by a dashed circle. **(b)** Schematic view on the sound generator’s lateral vibratory mass. **(c)** Schematic sagittal cross-section through the syrinx of the left bronchus, indicating the movement of pressures around and through the syrinx. Example raw data of sound production induced by bronchial and interclavicular air sac pressures **(d)** Example of fundamental frequencies produced in the pressure control space for the right hemi-syrinx of an adult male zebra finch (100 DPH). Phonation threshold pressures (PTP) are indicated for: PTP_icas_ (grey dot), and PTP_b_ (red dot). Transmural pressure (*p*_t)_ is indicated by white dashed lines for *p*_t_ = 0, 1 and 3 kPa. **(ef)** Frequency increases with *p*_t_ for both the left (dark red, dark blue) and right hemi-syrinx (light red, light blue) in males and **(c)** females, respectively. Black lines indicate linear regressions for region S1 (0 < *p*_t_ < 0.75) and S2 (*p*_t_ > 0.75). The phonation threshold pressure (PTP) values for individual *p*_icas_ ramps are indicated by black circles. **(g)** PTP for *p*_b_, *p*_t_, and *p*_icas_ (kPa), for the left and right hemi-syrinx. **(h)** Minimum frequency produced by the right hemi-syrinx was higher than the left hemi-syrinx (p = 0.09, see Table S1) for most individuals. **(i)** The absolute difference of minimal *f*_o_ was not significantly different between males and females. **(j)** Slopes for the left and right hemi-syrinx did not differ significantly between sides but were significantly steeper in S1 than in S2. For statistics see Supplementary Table S1. *, p<0.05; **, p<0.01; ***, p<0.001.

Tension and positioning of labia are modulated directly by two independent motor systems in songbirds: the respiratory and the syringeal motor system. The respiratory system generates the pressure and flow driving vibration of the avian vocal folds. In zebra finches, a sound pulse is produced when these tissues collide within the vibratory cycle^15^. Additionally, the syrinx is suspended in a pressurized air sac, the interclavicular air sac (ICAS), that is essential for sound production in birds^15,28,29^. The pressure differential between bronchial pressure (*p*_b_) and ICAS pressure (*p*_icas_) results in a net pressure, the transmural pressure (*p*_t_), that applies a net force on the MVM. This force has been shown to correlate most closely to changes in *f*_o_ during manipulation of *p*_icas_ *in vivo*^29^ and *ex vivo*^15^, but we do not know how these pressures interact to drive acoustic parameters.

The syringeal motor system of songbirds consists of up to 8 pairs of intrinsic and extrinsic syringeal muscles^21,30,31^ (Fig. 1a), which control the positioning of individual bones within the syringeal skeleton as well as the torque on them. Consequently, syrinx muscles thereby - either directly or indirectly – modulate the adduction level and tension of the labia, which are suspended between the bones^32^. The control space of syringeal muscles is vast due to its multidimensionality and only limitedly understood^15,31,33^. Taken together, both, changes in the motor control spaces or the sound production properties of the syrinx within this space could contribute to vocal development, but this is currently unknown.

Here we test if the acoustic output of the zebra finch syrinx sound generators changes over vocal development. We focused on the syringeal dynamics controlled by respiratory pressures and densely sampled the control space of bronchial versus air sac pressure.

## Materials and Methods

### *In vitro* sound production

#### Animals use and care

Zebra finches (*Taeniopygia guttata*, order Passeriformes) were kept and bred in group aviaries at the University of Southern Denmark, Odense, Denmark on a 12:12 hour light:dark photoperiod and given water and food *ad libitum*. All experiments were conducted in accordance with the Danish law concerning animal experiments and protocols were approved by the Danish Animal Experiments Inspectorate (Copenhagen, Denmark). Adult birds were provided with nesting material, *ad libitum*. Nest boxes were monitored daily, so that birds could be accurately aged. Breeding began in December 2017 and ended November 2018.

We studied vocal output of the syrinx *in vitro* in four age groups: 25 DPH (N = 5 males, N = 5 females), 50 DPH (N = 4 males, N = 4 females), 75 DPH (N = 6 males, N = 4 females), and 100 DPH (N = 5 males, N = 6 females). Sex was determined by dissection postmortem (25 DPH) or plumage (all other ages).

#### Syrinx mounting procedure

Animals were euthanized by Isoflurane (Baxter, Lillerød, Denmark) overdose. The syrinxes were dissected out while being regularly flushed and then submerged in a bath of oxygenated Ringer’s solution (recipe cf. ^34^) on ice. Each syrinx was photographed with a Leica DC425 camera mounted on a stereomicroscope (M165-FC, Leica Microsystems, Switzerland) prior to being placed inside the experimental chamber. Each bronchus was placed over polyethylene tubing (outer diameter 0.97 mm x inner diameter 0.58 mm, Instech Salomon, PA, USA) and secured with 10-0 nylon suture (AROSurgical, Newport Beach, CA, USA). The trachea was placed into a 1 mL pipette which was cut to the length of 15 mm (outer diameter 2.4 mm, inner diameter 1.4 mm). Some adipose tissue was left on the trachea to ensure an airtight connection. To control the pressure inside the chamber it was made airtight with a glass lid, which also allowed visualization of the syrinx through the stereomicroscope.

#### Experimental setup

We designed an experimental setup that allows studying the mechanical behavior of the syrinx *in vitro* or *ex vivo* (described in detail in^15^). The bronchial pressure (*p*_b_) and pressure in the experimental chamber (i.e. interclavicular air sac pressure, *p*_icas_) can be controlled independently by two dual-valve differential pressure PID controllers (model PCD, 0-10 kPa, Alicat Scientific, AZ, USA). Bronchial flow through the syrinx was measured with a MEMS flow sensor (PMF2102V, Posifa Microsystems, San Jose, USA) about 7 cm upstream from the bronchial connection. Sound was recorded with a ½ inch pressure microphone-pre-amplifier assembly (model 46AD, G.R.A.S, Denmark), amplified and high-pass filtered (0.2 Hz, 3-pole Butterworth filter, model 12AQ, G.R.A.S., Denmark). The sound recording sensitivity was tested before each experiment using a 1 kHz tone (sound calibrator model 42AB, G.R.A.S., Denmark). The microphone was placed 15 cm from the tracheal connector outlet and positioned at a 45° angle to avoid the air jet from the tracheal outlet. The sound, pressure and flow signals were low-pass filtered at 10 kHz (model EF120, Thor Labs) and digitized at 50 kHz (USB 6259, 16 bit, National Instruments, Austin, Texas).

#### Experimental protocol

To explore the acoustic output of the avian syrinx (Fig. 1a), we subjected the syrinxes to *p*_b_ ramps from 0-3 kPa (over atmospheric pressure) at 1 kPa/s with *p*_icas_ at a constant pressure. In one run, we applied 13 *p*_icas_ settings at equidistant 0.23 kPa intervals between 0 and 3 kPa in a randomized order with 2 s pause (*p*_b_ and *p*_icas_ = 0 kPa) between ramps (Fig. 1c). We avoided high flow regimes (*p*_t_ > 2 kPa) to decrease desiccation and avoid potential damage to the syrinx. When water in the bronchi or trachea prevented normal sound production in part of the run, we repeated the entire run. We subjected left and right hemi-syrinx separately in randomized order to this paradigm by allowing air flow only through one side and kept a one-minute rest between runs. The total duration for the experiment was maximally 20 minutes per individual.

#### Data Analysis

Sound was filtered with a 0.1 kHz 3^rd^ order Butterworth high-pass filter, pressure and flow signals were filtered with a 2 kHz 3^rd^ order Butterworth low-pass filter, all with zero-phase shift implementation (filtfilt function, Matlab). Sound, pressure and flow signals were binned in 2 ms duration bins, with a sliding window of 1 ms, and we calculated several parameters per bin. We calculated the RMS values for all signals. For the sound signal, we additionally extracted the fundamental frequency (*f*_o_), source level (SL) and Wiener entropy (WE).

Fundamental frequency was extracted using the yin-algorithm^35^ by manually optimizing minimal aperiodicity (0.05 – 0.2) and power (0.25 – 5.0 mPa). Source level at 1 m distance (SL) of the emitted sound was defined as:

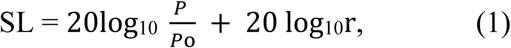

where *p* is the RMS of sound pressure (Pa), *p*_o_ is the reference pressure in air of 20 µPa, and r the distance (m) from the tracheal outlet to the microphone. To calculate WE, we first computed the amplitude spectrum Px of each bin using the periodogram method (nfft = 2048, overlap = 1024 samples). WE was defined as:

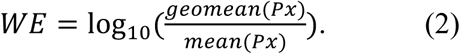

The transmural pressure over the MVM was defined as *p*_t_ = *p*_b_ -*p*_icas_. Phonation threshold pressures (PTP) were identified for *p*_icas_, *p*_b_, and *p*_t_ as the first pressure value for each *p*_b_ ramp where sound power crossed the set threshold. The minimal *f*_o_ was defined as the average *f*_o_ of the first value for each *p*_*b*_ ramp where sound power crossed the set threshold. Minimal SL was defined as the lowest SL value of all *p*_*b*_ ramps per side.

By combining pressure, flow and sound signals we were able to calculate the mechanical efficiency (ME) as the ratio (in dB) of acoustical power (*P*_acoustic_) over aerodynamic power (*P*_aerodynamic_):

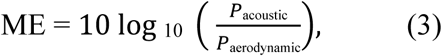

The acoustic power (W) was assumed to radiate over half a sphere: 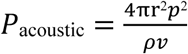, with air density *ρ* = 1.2 kg m^-3^, speed of sound *v* = 344 m s^-1^, and *p* the RMS sound pressure in Pa. The aerodynamic power *P*_*aerodynamic*_ = *p*_b_*V* where *p*_*b*_ is the bronchial pressure (Pa) and *V* is the flow rate (m^3^ s^-1^.) All data analysis was done in Matlab ^36^ and R version 3.5.1^37^.

#### MVM dimensions

After the experimental protocol, each syrinx was pinned down in the *situ* position and kept in 4% (w/v) paraformaldehyde for 24 hours at 4°C. It was then placed in PBS and stored at 4°C. To quantify the projected size of the MVM of each hemi-syrinx, we carefully cut the syrinx in half and made pictures of the medial side of the *medial tympaniform membranes* (MTMs) (Fig. 8d,e), and the lateral view of each hemi-syrinx using a Leica DC425 camera mounted on a stereomicroscope (M165-FC, Leica Microsystems, Switzerland) in Leica Application Suite (LAS Ver. 4.7.0). We measured the distances between physiological landmarks in the MTM following^20,32^: (i) the distance between the *medio-ventral cartilage* (MVC) and the *lateral dorsal cartilage* (LDC), and (ii) the distance between the LDC and the *medio-dorsal cartilage* (MDC). Distance measurements between cartilages were taken edge-to-edge in the middle of the cartilage. Additionally, we measured the area of the LDC which is fully suspended in the MVM. Length and area measurements were taken using ImageJ (Ver. 2.0.0-rc-69/1.52p;^38^).

### In vivo sound production

To measure the lowest fundamental frequency produced by an unactuated syrinx *in vivo*, we performed unilateral tracheosyringeal nerve lesions on the right side of the syrinx of juvenile and adult males.

#### Sound recordings

Song was recorded in custom-built, sound attenuated recording boxes (60×95×57 cm) and vocalizations were recorded continuously using a omnidirectional microphone (Behringer ECM8000) mounted 12 cm above the cage connected to an amplitude triggered recording software (Sound Analysis Pro;^39^). Sound was digitized at a sampling rate of 44.1kHz with 16-bit resolution (Roland octa capture, amplification 40 dB). Adult birds were recorded right before and 3 weeks after denervation. Juveniles were recorded at 50 and 100 DPH.

#### Tracheosyringeal nerve lesions

Unilateral denervations of the tracheosyringeal nerve were performed as previously described^40^. In brief, birds were anesthetized with isoflurane (induction 3%, maintenance 1.5 - 2%) and a 5 mm section of the right tracheosyringeal nerve was removed from the trachea through a small incision of the skin. After that the skin was re-sutured and the birds were allowed to recover in a heated cage. Adults were subsequently transferred to holding cages. Juveniles received surgery at 30 DPH and were returned to their parents after surgery.

#### Song analysis

Adult birds: For each bird (N = 5, age >120 DPH, days post-surgery: 20±2), the most common motif was determined by two experienced observers (IA, CPHE) in recordings acquired before denervation. Subsequently, the same motif was identified in recordings acquired 3 weeks after denervation. A custom written GUI (Matlab) was used to extract motifs from post denervation sound recordings. 200 motifs per bird were randomly selected for analysis. WAV files were high pass filtered at 0.3 kHz using a 3^rd^ order Butterworth filter. Syllables were assigned to be produced by the left or right hemi-syrinx based on whether their spectral content stayed the same or changed after denervation, respectively following earlier results by^41,42^.

Juvenile birds: Song of juvenile males (N = 6) was analyzed on recordings from 50 DPH (DPH 52.8 ± 1, days post-surgery: 22.3 ± 0.8). At this age, the syllable structure is not fully stereotyped, but motifs usually can be distinguished. We thus measured the lowest fundamental frequency of 50 motifs per bird irrespective of syllable identity. As the juvenile birds were denervated while singing subsong, we couldn’t distinguish which syllable was produced on which side. Under the assumption that the lowest fundamental is produced by the denervated side, we classified the measured syllables as right produced.

In all birds, frequencies were measured on spectrograms of post denervation recordings (FFT size: 1024, overlap: 75%, Hanning window) in SASlab (Avisoft, Germany). In all juvenile birds the frequency of the lowest harmonic (i.e *f*_o_) was measured. In all other cases, to increase the *f*_o_ resolution, we measured the 3 - 5th harmonic and *f*_o_ was calculated as:

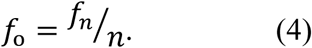

where *n* is the n^th^ harmonic. If several different syllables where analyzed for the adult birds the syllable with the lowest mean fundamental frequency (per side) was selected for statistical analysis.

### Statistics

To asses which pressure (*p*_b_, *p*_icas_, *p*_t_) drives *f*_o_ and SL in adults, we used a delta Bayesian information criterion (ΔBIC) approach. In a first step, we fit a linear model to the *f*_o_-pressure and SL-pressure relationship. We then computed the BIC values for these models and subtracted the BIC of the null model with no pressure variable (*f*_o_ = 1; SL = 1) to calculate the ΔBIC score. The model with the lowest ΔBIC score represents the model with the best fit.

To compare the *f*_o_ -*p*_t_ and SL – *p*_b_ relationship over ages and between side and sex, we fit individual linear models to the data and extracted the slopes. For *f*_o_ we split the data in two regions: region S1 (*p*_t_ = 0 – 0.75 kPa) and region S2 (*p*_t_ = 0.75 – 2 kPa).

To test whether age, sex or side of the syrinx had a significant effect on our response variables, we fit linear mixed effect models (LMMs) to our data using the lmer function of the lmerTest package^43^ with maximum likelihood optimization. Sex, side and age were fit as fixed effects and animal was included as a random effect to correct for dependence. Model equations can be found in Table S1, S2 and S3. To determine which effects significantly contribute to the model, we performed model selection using a Likelihood Ratio Test (LRT) with the Chi squared distribution using the lme4 package^44^. The difference between the absolute difference in minimal *f*_o_ between sexes and the difference in minimal *f*_o_ *in vivo* was assessed with Welch’s t-test. The difference between S1 and S2 slopes for *f*_o_ were assessed using paired t-tests for males and females.

All reported values in the text are mean ± SD. All error bars in figures are S.D. The outputs of al LMMs are reported in Supplementary Table S1, S2, and S3. We chose to present data in figures spilt by sex even when not significant. Statistical significance was accepted at p < 0.05 for all statistical tests.

## Results

We developed an experimental paradigm to quantify sound production as a function of driving parameters, because predicting the behavior of this complicated nonlinear system from anatomy or isolated mechanical property tests alone will only present correlates or rough estimates. Because we still have limited knowledge on the biomechanical effect of all syringeal muscles, we focused on syringeal dynamics controlled by respiratory pressures.

### Acoustic output of the adult syrinx

To quantify the functional acoustic output of the adult male and female zebra finch syrinx, we densely sampled the bronchial (*p*_b_) and interclavicular (*p*_icas_) pressure control space in isolated left and right hemi-syrinxes *in vitro*. For all zebra finch syrinxes tested, sound was produced in a pressure space enclosed by a minimal *p*_b_ and *p*_icas_ and exclusively in the lower half of the *p*_b_, *p*_icas_ space (Fig. 1d). Here a differential or transmural pressure (*p*_t_) exerts force on the MVM that is positive when directed outwards from bronchus to surrounding air sac. We quantified the phonation threshold pressures (PTP) for the three pressures (Fig. 1g). The PTP_b_ was not significantly different for sex (LMM, *p* = 0.418) or side (LMM, *p* = 0.395) and was 1.01 ± 0.28 kPa (range: 0.64 – 1.68 kPa, N = 11). The PTP_icas_ was also not significant different for sex (LMM, *p* = 0.638) or side (LMM, *p* = 0.223) and was 0.45 ± 0.27 kPa (range: 0.22-1.07 kPa, N = 11). The PTP_t_ was also not significantly different for sexes (LMM, *p* = 0.057) or side (LMM, *p* = 0.395) and was 0.15 ± 0.19 kPa (range: 4e-5 – 0.75 kPa, N = 11).

Next, we quantified the acoustic output of the adult zebra finch hemi-syrinx within the *p*_b_-*p*_icas_ control space for three parameters: fundamental frequency (*f*_o_), source level (SL) and Wiener entropy (WE). In all animals, fundamental frequency was gradually modulated depending on different combinations of *p*_b_ and *p*_icas_ (Fig. 1d-f). Frequency jumps were not observed. Of all pressures, *p*_t_ described the *f*_o_ data best (ΔBIC = −528, see Methods). The continuous, smooth increase of *f*_o_ with *p*_t_ can be clearly seen in Fig. 1e,f. In males, the right typically produced higher frequencies than the left as a function of *p*_t_ (Fig. 1e). However, the minimal *f*_o_ produced was not significantly different and was 511 ± 102 Hz (range: 352 – 613 Hz, N = 5) and 513 ± 123 Hz (range: 312 – 625 Hz, N = 5) for left and right hemi-syrinx respectively (Fig. 1h). To describe *f*_o_ as a function of *p*_t_, we split the data in to two regions (Fig. 1e,f); region S1 (*p*_t_ = 0 - 0.75 kPa) and region S2 (*p*_t_ = 0.75 - 2 kPa) and fit linear models to the data. The slope of S1 was 189 ± 76 Hz/kPa (range: 67 – 453 Hz/kPa, N = 5), and was significantly higher than the S2 slope of 55 ± 58 Hz/kPa (range: 25 – 310 Hz/kPa, N = 5) by 134 ± 18 Hz/kPa (Fig. 1j) (paired t-test, t = 5.3061, df = 9, *p* = 5e-4). In females, the minimal *f*_o_ produced was 529 ± 84 Hz (range: 435 – 678 Hz, N = 6) for the left and 568 ± 57 Hz (range: 518 – 650 Hz, N = 6) for the right hemi-syrinx (Fig. 1f). The slope of the S1 region was 181 ± 108 Hz/kPa (range: 67 – 453 Hz/kPa, N = 6) and was also significantly higher than the S2 slope of 55 ± 58 Hz/Pa (range: −42 – 167 Hz/kPa, N = 6) by 126 ± 51 Hz/kPa (Fig. 1j) (paired t-test, t = 3.31, df = 10, *p* = 0.0079). Comparing the sexes, the difference in minimal *f*_o_ between left and right seemed smaller in males than in females (Fig. 1i), but this effect was not significant (Welch’s t-test, t = −1.08, df = 6.42, p = 0.32). The slopes of both S1 and S2 were not significant different between sex and side and were 185 ± 92 Hz/kPa (range: 25 – 453 Hz/kPa, N = 11) and 55 ± 58 Hz/kPa (range: −42 - 167, N = 11), for S1 and S2, respectively (Fig. 1j, Supplementary Table S1).

Source level at 1 m distance was best described by *p*_b_ (ΔBIC = −78,996) and increased linearly with pressure in both sexes (Fig. 2a-c). The minimum SL (Fig. 2d) was not significantly different for sex (LMM, *p* = 0.149) or side (LMM, *p* = 0.225) and was 45 ± 4 dB re 20µPa @ 1m (range: 37 - 51 dB re 20µPa @ 1m, N = 11). The slope of the SL-*p*_b_ relationship (Fig. 2e) did not differ significantly between males and females (LMM, *p* = 0.704), but was significantly higher on the right side (5 ± 2 dB/kPa, N = 11) than the left side (4 ± 1 dB/kPa, N = 11) (LMM, *p* = 0.016).

**Figure 2.**
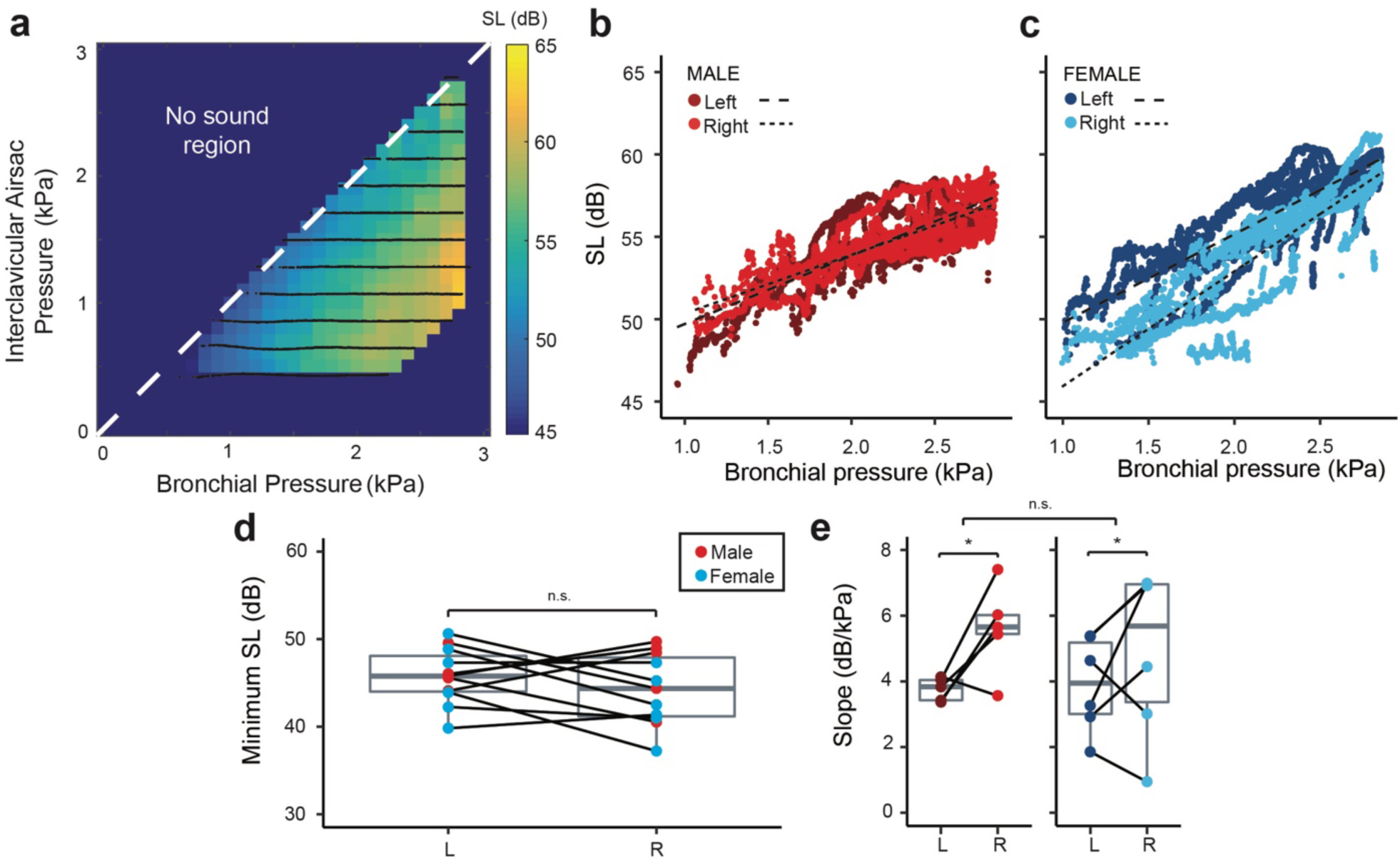
Source level increases with bronchial pressure in the adult zebra finch hemi-syrinx. **(a)** Example of source level (SL) produced in the pressure control space for the left hemi-syrinx of an adult male zebra finch (100 DPH). Source level increases with *p*_*b*_ for both the left (dark red, dark blue) and right hemi-syrinx (light red, light blue) in **(b)** males and **(c)** females, respectively. Black dashed lines indicate linear regressions (Left: long dash; Right: short dash). **(d)** Minimum SL does not differ significantly between the left and right hemi-syrinx or between males and females. **(e)** The SL increase with bronchial pressure was significantly steeper for the right hemi-syrinx than the left hemi-syrinx. For statistics see Supplementary Table S1. *, p<0.05.

WE did not show any obvious relation to *p*_b,_ *p*_icas_, or *p*_t_, and therefore we considered the mean of the entire control space (Fig. 3a). The mean WE did not differ significantly between sexes (LMM, *p* = 0.138) or sides (LMM, *p* = 0.661) and was −1.8 ± 0.12 dB (range: −1.6 to −2.1 dB, N = 11) (Fig. 3b).

**Figure 3.**
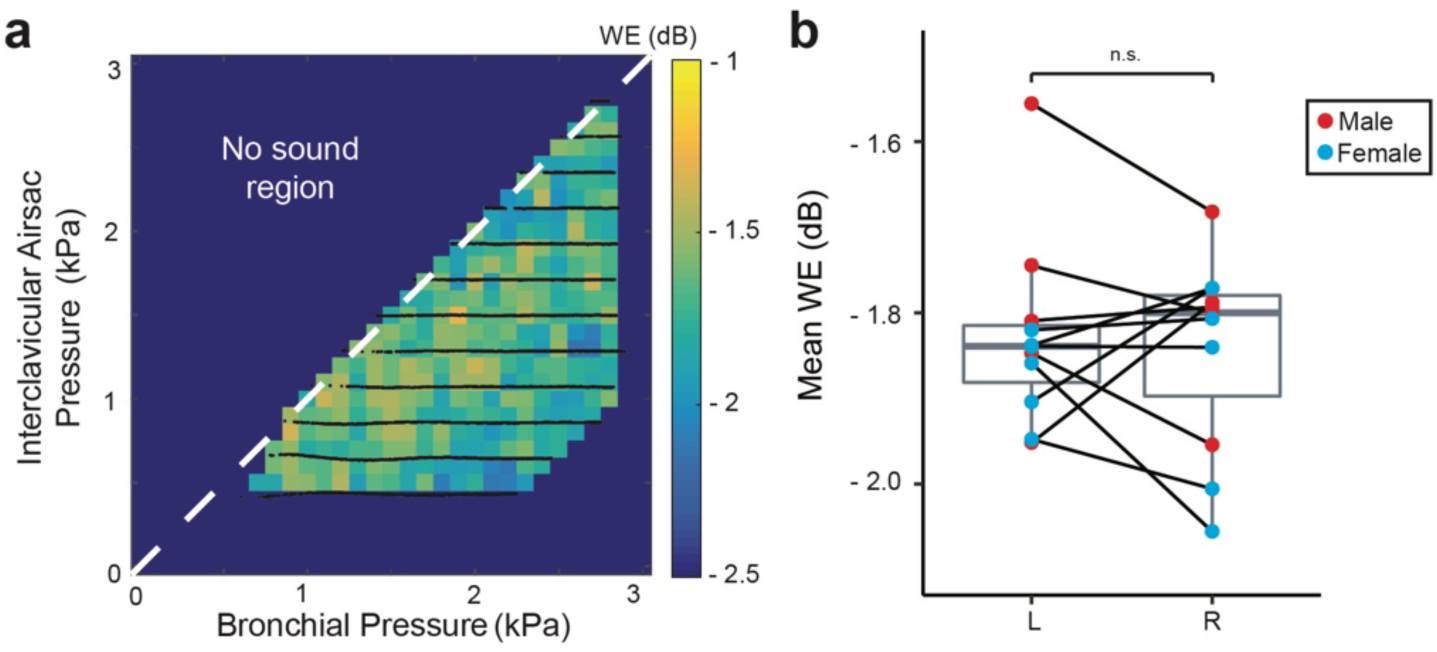
Wiener entropy is not affected by pressure in the adult zebra finch hemi-syrinx. **(a)** Wiener entropy (WE) produced in the pressure control space for the left hemi-syrinx of an adult male zebra finch (100 DPH). **(b)** Mean WE of the left and right hemi-syrinx was not significantly different between male and female zebra finches or for left and right hemi-syrinxes. For statistics see Supplementary Table S1.

Our setup allowed us to calculate the mechanical efficiency (ME) of sound production, which estimates how much of the power in the air flow is transformed into sound (see Methods, Fig. 4). We found that ME did not vary systematically with *p*_b,_ *p*_icas_, or *p*_t_ and therefore considered the mean of the entire control space (Fig. 4b). ME was not significantly different between sexes (LMM, *p* = 0.118), but was significantly lower (LMM, *p* = 0.027) on the right side (−36 ± 1.8 dB, range: −39 to −33, N = 10) compared to the left (−35 ± 2.2 dB, range; −38 to −31 dB, N = 10) (Fig. 4c).

**Figure 4.**
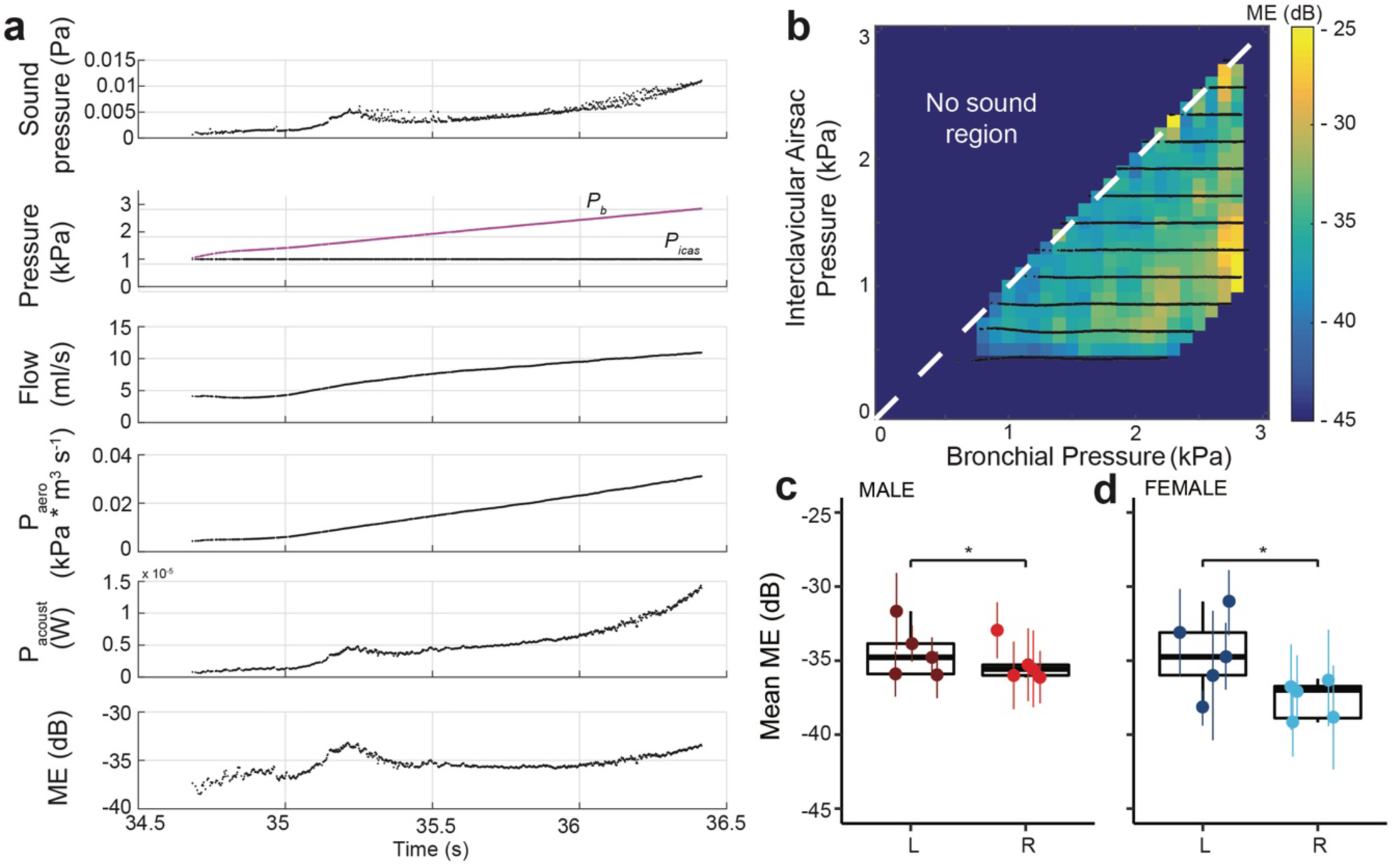
Mechanical efficiency is not affected by respiratory pressure, but lower on the right side of adult zebra finch. **(a)** Example of one *p*_b_-ramp (from top to bottom) received sound pressure (Pa), bronchial and interclavicular air sac pressures (kPa), acoustic and aerodynamic power (W), and mechanical efficiency (ME) (dB). **(b)** Mechanical efficiency in the pressure control space for the left hemi-syrinx of an adult male zebra finch (100 DPH). **(c, d)** Mechanical efficiency was significantly higher for the left hemi-syrinx than the right hemi-syrinx in both sexes. For statistics see Supplementary Table S1. *, p<0.05.

#### Acoustic control space does not change over vocal development

Next, we tested if the output of the isolated zebra finch syrinx changed over vocal development from 25 to 100 DPH (Fig. 5). Like in the adult syrinx, sound was produced exclusively in the lower half of the syringeal *p*_b_, *p*_icas_ space over vocal development. The PTP_b_ was not significantly affected by age (LMM, *p* = 0.257), side (LMM, *p* = 0.268), or sex (LMM, *p* = 0.558) and was 0.9 ± 0.24 kPa (range: 0.53 – 1.68 kPa, N = 32). PTP_icas_ demonstrated a small but significant increase with age (LMM, *p* = 0.002), but no significant difference between sexes (LMM, *p* = 0.762) or sides (LMM, *p* = 0.274), and went from 0.21 ± 0.8 kPa at 25 DPH (range: 0.006 – 0.25 kPa, N = 8) to 0.37 ± 0.25 kPa at 100 DPH (range: 0.21 – 1.1 kPa, N = 11).

**Figure 5.**
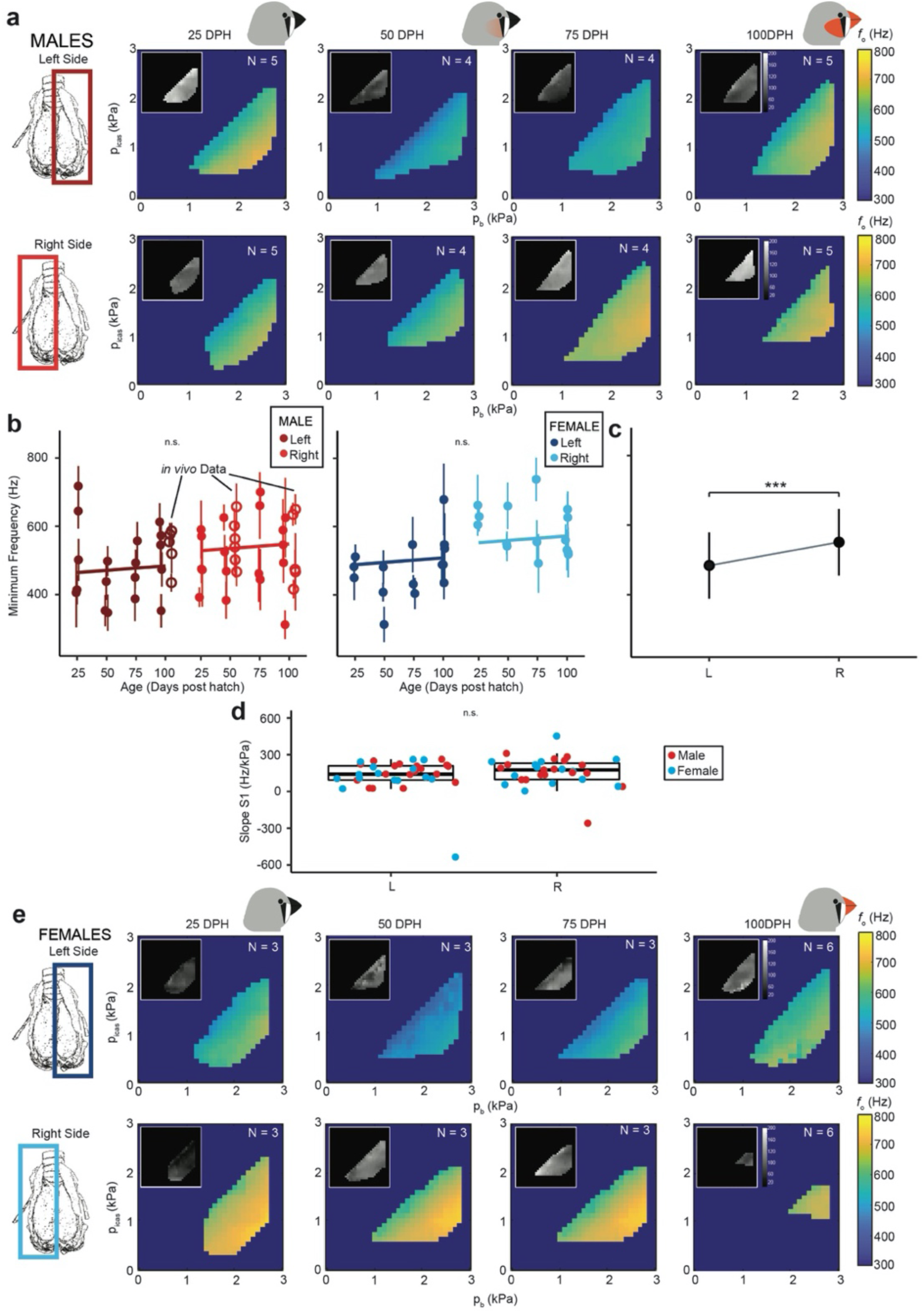
Minimal frequency does not change over zebra finch song development. **(a)** Fundamental frequency in the pb picas pressure control space for the left and right hemi-syrinx in males over vocal development *f*_o_ values are averaged over all individuals within a group (insets S.D.). **(b)** Minimum *f*_o_ does not change with age in males and females. **(c)** Minimum frequency was significantly different between the left and right hemi-syrinx, regardless of age or sex. **(d)** Slope S1 for the relationship between *f*_o_ and *p*_t_ was not significantly different for side. **(e)** Pressure control spaces *in vitro* for the left and right hemi-syrinx in females. For statistics see Supplementary Table S2. ***, p<0.001.

Consistently, PTP_t_ also demonstrated a small but significant decrease with age from 0.17 ± 0.08 kPa at 25 DPH (range: 0.06 – 0.31 kPa, N = 8) to 0.03 ± 0.06 kPa at 100 DPH (range: 4e-5– 0.16 kPa, N = 11), and marginally, but significantly lower in males (0.0846 ± 0.0981 kPa) compared to females (0.1047 ± 0.0875).

We examined changes in the syringeal *p*_b_-*p*_icas_ control space for three acoustic parameters: fundamental frequency (*f*_o_), source level (SL) and Wiener entropy (WE). Over development, *f*_o_ was also gradually modulated within the *p*_b-_*p*_icas_ space and did not exhibit frequency jumps (Fig. 5). In males and females, the minimal *f*_o_ produced by each hemi-syrinx *in vitro* did not change significantly from 25 DPH to 100 DPH (Fig. 5b), but the left hemi-syrinx produced a significantly lower minimal *f*_o_ than the right (males: L: 487 ± 106 Hz, N = 18, R: 519 ± 104 Hz, N = 18, females: L: 480 ± 83 Hz, N = 15, R: 590 ± 69 Hz N = 15) (LMM, *p* < 0.001). Interestingly, in 5 out of 18 males the left-right difference was reversed.

The slope of the *f*_o_-*p*_t_ relationship in region S1 did not significantly change with age (LMM, *p* = 0.651), sex (LMM, *p* = 0.630), or side (LMM, *p* = 0.270) and was 91 ± 109 Hz/kPa (range: −204 – 250 Hz/kPa, N = 32). The slope in region S2 was significantly different between ages (LMM, *p* = 0.0273), but not for sex (LMM, *p* = 0.464), or side (LMM, *p* = 0.474), and was 88 ± 43 Hz/kPa (range: 31 – 146 Hz/kPa, N = 8), 62 ± 28 Hz/kPa (range: 18 – 105 Hz/kPa, N = 7), 70 ± 46 Hz/kPa (range: 5 – 149 Hz/kPa, N = 7), and 65 ± 50 Hz/kPa (range: −43 - 135, N = 11) for 25, 50, 75, and 100 DPH respectively.

Source level increased with *p*_b_ for all animals (Fig. 6a,e), but the minimal source level did not change significantly with age (LMM, *p* = 0.680), sex (LMM, *p* = 0.374), or side (LMM, *p* = 0.271) and was 44 ± 4 dB re20µPa @ 1m (range: 36 – 51 dB, N = 32) (Fig. 6b-c). The SL-*p*_b_ slope also did not change with age (LMM, *p* = 0.83), or sex (LMM, *p* = 0.874), but was significantly less steep (LMM, *p* = 0.037) on the left (4.1 ± 1.0 dB/kPa, range: 1.9 – 6.2 dB/kPa, N = 32) versus right hemi-syrinx (4.7 ± 1.6 dB/kPa, range: 0.9 – 7.4 dB/kPa, N = 32) by 0.6 ± 0.3 dB (N = 32).(Fig. 6d). Mean WE did not change significantly for any of the parameters tested (LMM, age: *p* = 0.070, sex: *p* = 0.979, side: *p* = 0.084), and was −1.8 ± 0.1 dB (range: - 2.1 to −1.6 dB, N = 32; Fig. 7a-b, e-f).

**Figure 6.**
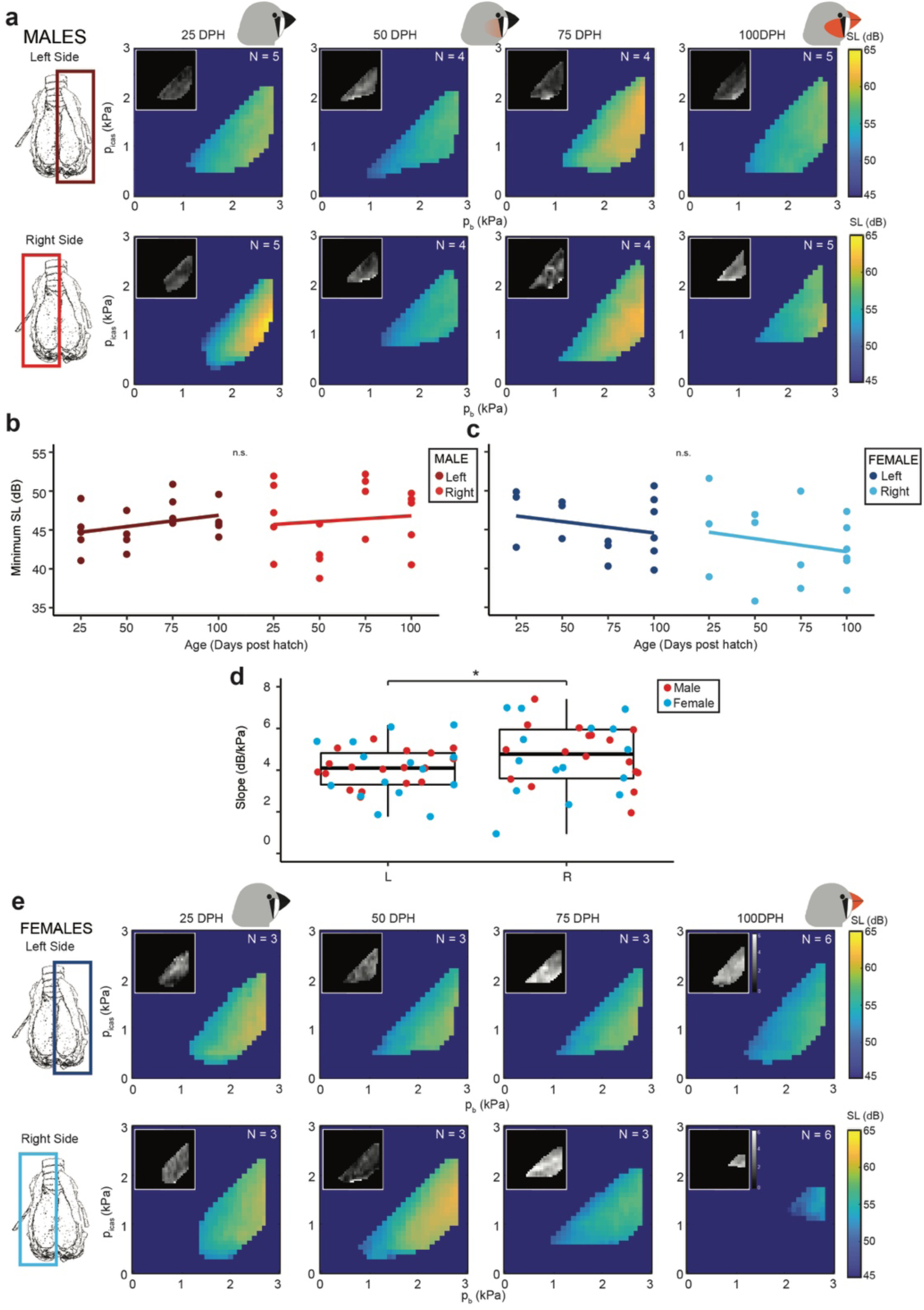
Minimum source level (SL) does not change over zebra finch song development. **(a)** Mean SL pressure control spaces *in vitro* for the left and right hemi-syrinx in males (inset S.D.). Minimum SL produced for **(b)** males and **(c)** females does not change over development, nor was it significantly different between sides. **(d)** Slope for the relationship between SL and *p*_b_ was significantly steeper on the right compared to the left hemi-syrinx. **(e)** Mean SL pressure control spaces *in vitro* for the left and right hemi-syrinx in females. For statistics see Supplementary Table S2. *, p<0.05.

**Figure 7.**
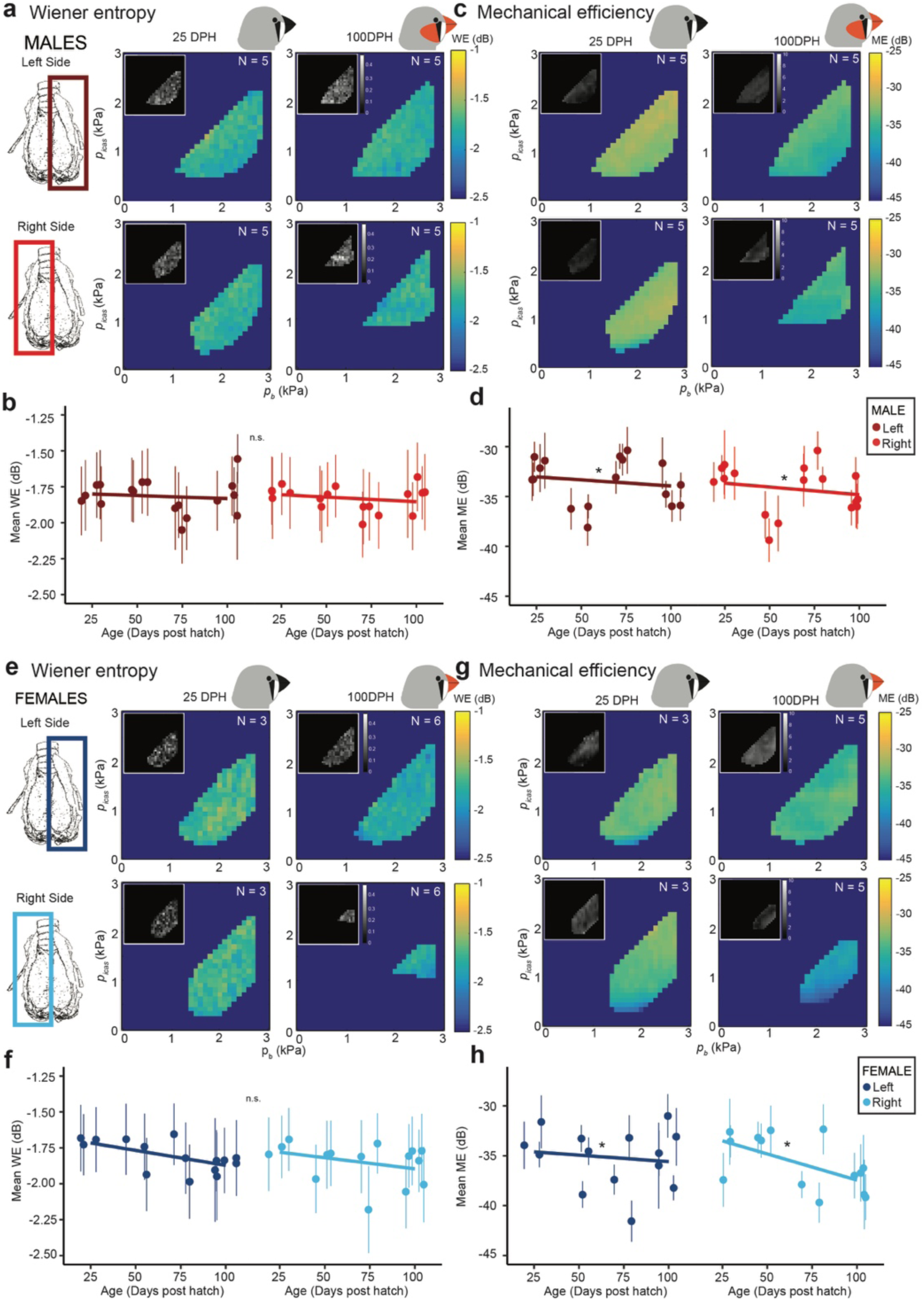
Wiener entropy and mechanical efficiency do not change over zebra finch song development. **(a)** Wiener entropy (WE) for the left and right hemi-syrinx in males age 25 and 100 DPH. **(b)** WE did not change for males over development. **(c)** Mechanical efficiency (ME) for the left and right hemi-syrinx in males age 25 and 100 DPH. **(d)** The mean ME produced for males significantly differed over development. **(e)** Wiener entropy (WE) for the left and right hemi-syrinx in females age 25 and 100 DPH. **(f) In females**, WE did not significantly change over development. **(g)** Mechanical efficiency (ME) for the left and right hemi-syrinx in females age 25 and 100 DPH. **(h)** The mean ME of female syrinxes decreased significantly over development. For statistics see Supplementary Table S2. *, p<0.05.

Next, we calculated the mechanical efficiency (ME) to test if the ability of the syrinx to convert mechanical power to sound changed over song development (Fig. 7c,g). ME did not significantly change with sex (LMM, *p* = 0.091) or side (LMM, *p* = 0.101), but demonstrated a small but significant decrease (LMM, *p* = 0.046) from −33 ± 1.7 dB (range: −37 to −31 dB, N = 8) at 25 DPH to −35 ± 1.8 dB (range: −39 to −33 dB, N = 10) in adults (Fig. 7d, h).

Taken together, we observed that the acoustic output of the syrinx control space did not change considerably over development in males and females *in vitro*. To test whether these *in vitro* results are representative for the performance of the syrinx during *in vivo* sound production, we measured the acoustic output over development *in vivo*. Because both SL and WE are affected by vocal tract filtering properties, we focused on fundamental frequency. We measured the lowest fundamental frequency produced by males with unilateral syrinx denervation at 50 and 100 DPH (see Methods). Consistent with the *in vitro* data, the minimal *f*_o_ did not change significantly (Welch’s t-test, t = 0.49, df = 6.60, p = 0.64) with age (Right hemi-syrinx: 555 ± 70 Hz (N = 6) and 524 ± 98 Hz (N = 5) for 50 and 100 DPH respectively. Left adult hemi-syrinx: 516 ± 54 Hz (N = 5) and also did not differ between the left (intact) and right (denervated) hemi-syrinx. Furthermore, the *in vivo* values were consistent with the *in vitro* data (Fig. 5b), strongly suggesting that the output of the syrinx *in vitro* is representative for the *in vivo* situation.

#### Morphological changes over development

We lastly investigated if the medial vibratory mass (MVM) dimensions change over development. We focused on non-invasive photographic techniques because detailed histological or material property testing was beyond the scope of the present study. The tension in the MVM can be increased by the shortening of two syringeal muscles that insert on the *medio-ventral cartilage* (MVC) and *medio-dorsal cartilage* (MDC)^21^ (Fig. 8a-c). A third cartilage, the *lateral dorsal cartilage* (LDC), is embedded within the MVM. A thickening of the tissue between the MVC and LDC is called the medial labium. The two syringeal muscles change the tension of the MVM by changing the distance between 1) MVC and LDC and 2) LDC and MDC (Fig. 8b,c). The MVC - LDC length did not change with sex (LMM, *p* = 0.72) or age (LMM, *p* = 0.446), but was significantly (LMM, *p* = 0.002) longer on the left (1.13 ± 0.10 mm; range: 965 – 1297 μm, N = 29) compared to the right (1.05 ± 0.11 mm, range: 798 – 1363 μm, N = 29) (Fig. 8f). The LDC - MDC length also did not change with sex (LMM, *p* = 0.982) or age (LMM, *p* = 0.191), but it was significantly (LMM, *p* = 0.149) longer on the left (0.71 ± 0.16 mm, range: 387 – 1107 μm, N = 29) compared to right (0.65 ± 0.13 mm, range: 365 – 858 μm, N = 29) (Fig. 8g). The area of the LDC was smaller (LMM, *p* < 0.001) on the left (4.1e4 ± 2.2e4 μm^2^, range: 1978 – 9.9e4 μm^2^, N = 29) compared to the right (6.2e4 ± 3.0e4 μm^2^, range: 11867 – 1.1e5 μm^2^, N = 29) hemi-syrinx and increased over development (LMM, *p* = 0.048; Fig. 8h). Lastly, we approximated overall growth of the syrinx as the distance between both ends of the B4 cartilage in the bronchus, and found no significant change for side (LMM, *p* = 0.14), sex (LMM, *p* = 0.17), or age (LMM, *p* = 0.8) and was 1.85 ± 0.23 mm (range: 1280 – 2196 μm, N = 29; Fig. 8i).

**Figure 8.**
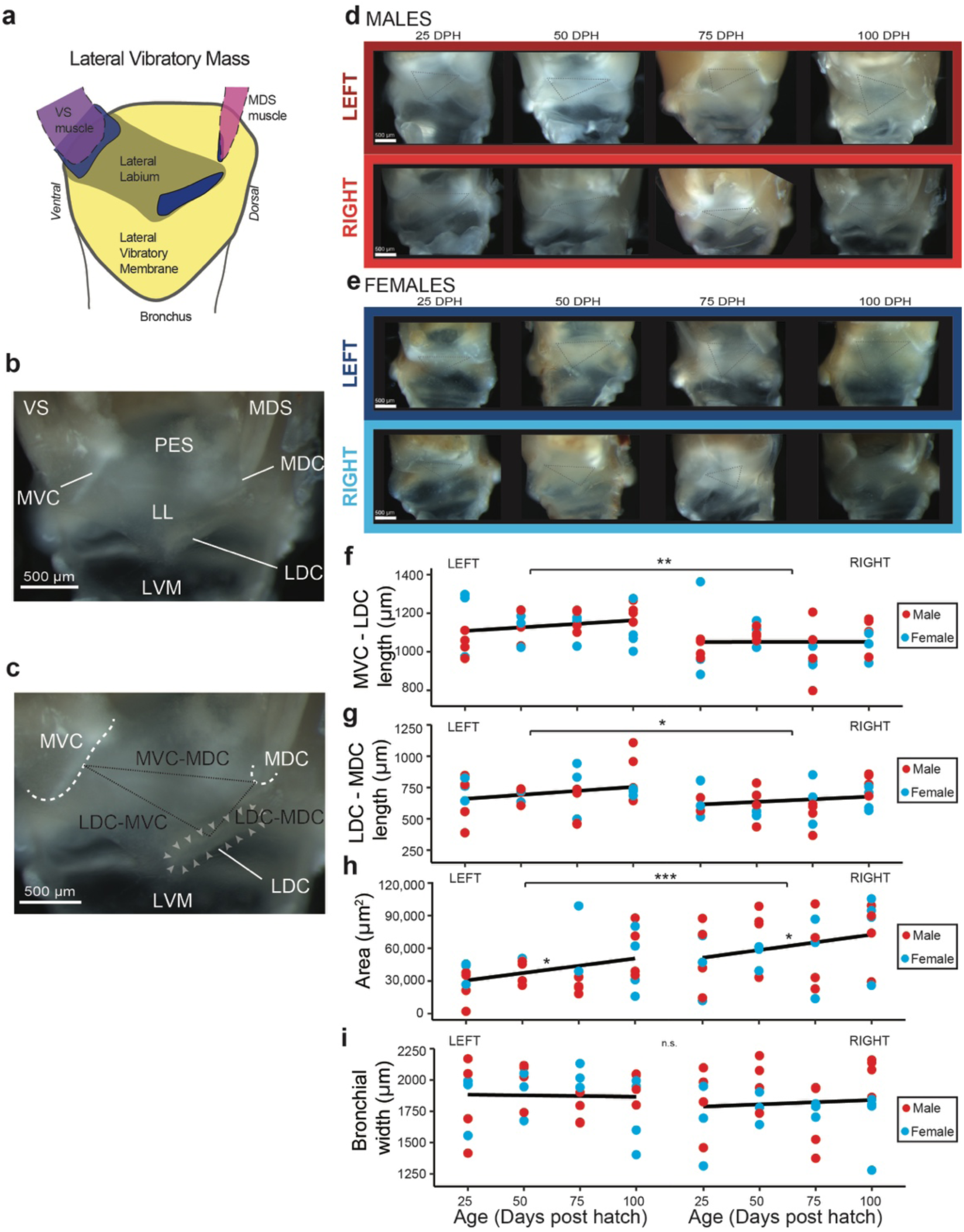
Morphological changes of the medial labia over zebra finch song development. **(a)** Schematic of lateral vibratory mass. **(b)** Physiological landmarks in the ML, the *medio-ventral cartilage* (MVC), the *medio-dorsal cartilage* (MDC) and the *lateral dorsal cartilage* (LDC) and **(c)** the distance between landmarks. **(d**,**e)** Example images of MTMs of left/right and male/female hemi-syrinx. **(f)** MVC-LDC length was significantly different between sides, but not over development. **(g)** LDC-MDC length significantly differed between sides, but not over development. **(h)** LDC area for the four investigated age groups was significantly different and differed significantly between the two sides. **(i)** Bronchial width did not significantly differ over development or side. For statistics see Supplementary Table S3. *, p<0.05; **, p<0.01; ***, p<0.001.

## Discussion

As songbirds learn to sing, both their song system and vocal organ are undergoing postnatal changes that lead to dramatic changes in vocal behavior. Here we show that the acoustic output of the sound generators within the zebra finch syrinx does not change considerably over vocal development. This strongly suggests that the observed acoustic changes during vocal development are caused by changes in the motor control pathway, from song system circuitry to muscle force, and not by changes in the avian analog of the vocal folds. In other words, the size and properties of the instrument are not changing, but its player is.

We show that both sides of the syrinx, or hemi-syrinxes, produce sound in a well-defined consistent pressure control space. The pressure space is bound by phonation threshold pressures (PTP) for *p*_b_, *p*_icas_ and *p*_t_, which are the same for both sexes and sides. The PTP_b_ values in adults of 1.01 ± 0.28 kPa reported here are consistent with earlier reported PTP_b_ values of 1.2 kPa in situ^45^ and *in vivo*^31^. The possibility to control the two respiratory pressures (*p*_b_ and *p*_icas_) independently is very different from laryngeal systems where no air sacs directly apply force on the vocal folds. To what extent birds can actively and independently control the magnitude of *p*_t_ *in vivo* during song remains unknown. Certainly, positive pressurization of *p*_icas_ is an essential condition to achieve syringeal vibration in all investigated species *in vivo*, (e.g., zebra finches^29^ and *in vitro*^15,46,47^, and dynamic *p*_t_ changes up to 1 kPa occur during vocalizations in ringdoves^34,48–50^.

Our data show that source level and fundamental frequency are modulated by respiratory pressures. Source level was driven by bronchial pressure, consistent with laryngeal voiced sound production^19^. Fundamental frequency (*f*_o_) was set predominantly by *p*_t_, consistent with the idea that *p*_t_ acts as a force on the MVM that increases tension^51,52^, which has also been observed in collapsible tubes^54^. Interestingly, especially close to the phonation onset *f*_o_ modulation is steep (200 Hz/kPa) and a dynamic *p*_t_ difference of 1 kPa could in zebra finches lead to *f*_o_ modulation of comparable magnitude to full stimulation of the ventral syringeal muscle^15^. In contrast, Wiener Entropy (WE) was not modulated systematically in the pressure space. However, over the course of song learning WE has been shown to change over multiple time scales, from within a day to weeks and months^2,3,55^ and increased entropy variance is linked to better learning success^3,55^. These data suggest that WE is under control of the motor systems of (i) the syrinx, by modulating collision force and thereby spectral slope of the sound source^56^, and (ii) the upper vocal tract, by changing resonance properties and thereby frequency content of the radiated sound^57,58^.

Mechanical efficiency may have been an important selecting factor during the evolution of the syrinx as a vocal organ^59^ and our data confirms that the mechanical efficiency of voiced sound production is indeed high in the avian syrinx compared to the mammalian larynx. Our data shows that about 0.05% (−33 dB) of the flow energy is converted into sound, while in mammals, laryngeal ME is only about 1.10^−3^ to 0.01% (−50 to −40 dB) in tigers (Titze et al., 2010) and 3.10^−5^ to 1.10^−3^ % (−65 to −50 dB) in marmosets^6^). ME in the rooster has been reported to be as high as 1.6% *in vivo*^61^. The ME of the zebra finch *in vivo* is likely to be higher than we measured *in vitro*, because our preparation does not include a vocal tract. The reported SL of the zebra finch is about 70-80 dB re 20µPa @ 1m^62^, while the SL of our isolated syrinx was 20-30 dB SPL lower. A conservative estimate of 10 dB SL increase (i.e. RMS is 3 times higher) while maintaining the same aerodynamic power, results in a 10-fold increase in radiated acoustic power and a ME of 0.5% (−23 dB). This is already comparable to the very high efficiency reported by Brackenbury^61^. Furthermore, we observed slight differences of ME over development, but these effects were not as pronounced as the changes observed in marmosets, where the adult larynx produces louder and more efficient vocalisations than the infant larynx^6^. Thus, the ME of the zebra finch syrinx is very high already at 25 DPH and remains unaffected by vocal development.

Our data shows that acoustic output of the syrinx is modulated smoothly within the pressure boundaries: both SL and *f*_o_ increased continuously with bronchial and air sac pressures. Thus when driven by physiologically relevant pressures, this nonlinear dynamic system exhibits only one bifurcation from steady (no vibration and no sound) to flow-induced self-sustained vibration, without additional bifurcations, such as period doublings or jumps to deterministic chaos^45,63–66^. Earlier work already showed that in a subset of the pressure control space *in situ*, additional bifurcations did not occur^45^. Moreover, the supposed bifurcations during song *in vivo*, e.g. frequency jumps, were shown to not be actual bifurcations^45^, but more likely millisecond scale modulation driven by superfast syringeal muscle^67^. Here we also show that in the independently controlled and densely sampled pressure control space of the zebra finch syrinx *in vitro*, additional bifurcations do not occur. Thus, driven by physiologically relevant pressures, the acoustic output of the syrinx is continuous, which simplifies sensory-motor control^45^.

The stable acoustic output of the sound generators over development suggests that its mechanical properties are not changing^22,23^ and acoustic output^41,68,69^ are common in songbirds. In all investigated species to date, except for the Bengalese finch^15,69^, the left hemi-syrinx produces lower frequencies.

The *f*_o_ of sound is determined by positioning of and driving pressures on the vocal folds, as well as their resonance properties^20^. Small changes in material properties can drive changes in vocalizations over development in marmosets^6^. Because we show that minimum frequency does not significantly change over development *in vitro* and *in vivo*, our data strongly suggest that the effective resonance properties of the syringeal sound generators when driven by physiological realistic pressures do not change over vocal development. Next to resonance properties, the change in MVM stiffness as response to shortening is also likely changing very little over development. Although we didn’t measure directly how syringeal muscle force modulates MVM resonance properties over development, the *f*_o_-*p*_t_ relationship (S1) can be seen as a proxy of the effect of MVM lengthening by e.g. VS contraction, as *p*_t_ exerts a force on the MVM that causes changes in strain^32^. Taken together, the MVM vibration frequencies and its response to external force do not change, which strongly suggest that its material properties also do not change over development.

We propose that syringeal sound production on the one hand, and its control on the other, are shaped differentially by genetic and environmental factors. First, we propose that the development and maintenance of the sound generators is predominantly under genetic control. Second, we propose that the syringeal muscles and skeletal properties are heavily influenced by use, which can ultimately lead to a sexually dimorph syrinx.

Our data suggest that MVM mechanical properties are set and maintained by genetic factors. First, a salient feature in our dataset is that the adult left hemi-syrinx produces a lower minimal *f*_o_ compared to the right in both sexes, corroborating earlier findings^20^. In most songbirds both hemi-syrinxes contribute to the vocal repertoire^70,71^ and left-right differences in labial morphology^22,23^ and acoustic output^41,68,69^ are common in songbirds. In all investigated species to date, except for the Bengalese finch^15,69^, the left hemi-syrinx produces lower frequencies. Here we show for the first time that differences between left and right sound generators, size and acoustic output, are already established at 25 DPH. Second, we show that neither the morphology, nor the acoustic output of the bilateral sound generators changes over development. Thus, the MVM properties have matured already at 25 DPH. This latter feature makes the songbird system a wonderful model system to study vocal development compared to mammalian systems where larynx maturation strongly contributes to vocal output^6,25–27^. Third, MVM properties remain constant even though they are colliding billions of times over their lifetime and there must be a large variability in use between individuals and sexes. In human vocal folds, strong collisions can lead to several types of lesions that severely affect vocal output^72^. Thus, ultrastructural repair of vibratory tissues must lead to maintenance of their properties, but what cellular and molecular mechanisms underlie this maintenance in birds remains unknown. Taken together, these observations support the ideas that the sound generators of the syrinx are matured at the onset of vocal learning, their composition remains stable and is species specific, which suggests that their composition and dynamical behavior is set by genetic factors and not affected by use. The zebra finch syrinx is thus a stable instrument which’s postnatal development does not add further complexity to song learning

In stark contrast, syringeal muscles are changing functionally over postnatal development during song learning. Until 25 DPH the syrinx of male and female zebra finches is not clearly distinguishable. However, after 25 DPH, the syringeal muscles start to exhibit sexually dimorphic features, such as increased muscle mass^11^ and speed in males^12^, which leads to a different action potential to force transform^14^. Furthermore, muscle force exerted on bones is known to redirect bone deposition and thus can lead to significant bone remodelling^73^. Taken together, we propose that in songbirds syringeal muscle activity and resulting forces acting on the syrinx are crucially important drivers to the sexual dimorphism of bones and cartilages of the syringeal skeleton. This implies that most anatomical changes are caused by muscle use and training driven by the sexually dimorphic songbird brain (i.e., song system). As such the instrument, i.e. the vocal folds, are not changing, but the player is.

## Acknowledgments

The authors wish to thank Torben Christensen and Sonja Jakobsen for technical support, Emil B. Hansen for help with breeding, and John Jackson for assistance with statistical analysis.

## Author Contributions

A.M., I.A. & C.E. conceived the study. A.M., I.A. & C.E. developed technical methods. I.A. & C.E. provided reagents and materials. A.M. & P.L. performed *in vitro* experiments and collected all data. I.A., P.S. & C.E. performed *in vivo* experiments and collected all data. A.M., I.A. & C.E. analysed the data. A.M. prepared all figures and tables. A.M., I.A. & C.E. wrote the manuscript. All authors read and approved the final version of the manuscript.

## Additional Information

The authors declare that they have no competing interests.

## Funding

This work was supported by the Danish Research Council (Grant DFF 5051-00195), the Carlsberg Foundation (Grant CF17-0949) to I.A. and Novo Nordisk Foundation (Grant NNF17OC0028928) to C.P.H.E

## Data Availability

The datasets generated during and/or analysed during the current study are available from the corresponding author on reasonable request.

## Notes

### Competing Interest Statement

The authors have declared no competing interest.

