## Supplementary Tables for "Syringeal vocal folds do not have a voice in zebra finch vocal development"

**Table S1. Linear mixed effects model outputs for the acoustic results in adult zebra finches.**

| | Model parameter | Estimate of slope | Std. error | F value | DF | p |
|---|---|---|---|---|---|---|
| | $PTP_b = \beta_0 + \beta_{side} + \beta_{sex} + \epsilon$ | | | | | |
| **$PTP_b$** | Intercept | 1.0 | 0.10 | | | <0.001 |
| | Sex | -0.1 | 0.12 | 0.69 | 1 | 0.418 |
| | Side | 0.1 | 0.12 | 0.76 | 1 | 0.395 |
| | $PTP_b = \beta_0 + \beta_{side} + \beta_{sex} + \epsilon$ | | | | | |
| **$PTP_t$** | Intercept | 0.2 | 0.06 | | | 0.007 |
| | Sex | -0.2 | 0.07 | 4.11 | 1 | 0.057 |
| | Side | 0.1 | 0.07 | 0.76 | 1 | 0.395 |
| | $PTP_{icas} = \beta_0 + \beta_{side} + \beta_{sex} + \epsilon$ | | | | | |
| **$PTP_{icas}$** | Intercept | 0.4 | 0.10 | | | 0.002 |
| | Sex | 0.1 | 0.11 | 0.23 | 1 | 0.638 |
| | Side | 0.1 | 0.11 | 1.59 | 1 | 0.223 |

| | Model parameter | Estimate of slope | Std. error | LL $\chi^2$ | DF | p |
|---|---|---|---|---|---|---|
| | $Minf_o = \beta_0 + \beta_{side} + \beta_{sex} + \epsilon$ | | | | | |
| **Min $f_o$** | Intercept | 539.6 | 24.40 | | | <0.001 |
| | Sex | -34.0 | 35.10 | 0.90 | 1 | 0.34 |
| | Side | 21.1 | 12.40 | 2.85 | 1 | 0.09 |
| | $S1f_o = \beta_0 + \beta_{side} + \beta_{sex} + \epsilon$ | | | | | |
| **S1 $f_o$** | Intercept | 172.7 | 32.62 | | | <0.001 |
| | Sex | 17.6 | 37.59 | 0.03 | 1 | 0.859 |
| | Side | 7.2 | 40.49 | 0.22 | 1 | 0.640 |
| | $S2f_o = \beta_0 + \beta_{side} + \beta_{sex} + \epsilon$ | | | | | |
| **S2 $f_o$** | Intercept | 69.0 | 19.82 | | | 0.0021 |
| | Sex | -17.2 | 23.55 | 0.27 | 1 | 0.606 |
| | Side | -12.3 | 23.65 | 0.53 | 1 | 0.468 |

| | | | | | | |
|---|---|---|---|---|---|---|
| | $MinSL = \beta_0 + \beta_{side} + \beta_{sex} + \epsilon$ | | | | | |
| **Min SL** | Intercept | 44.7 | 1.22 | | | <0.001 |
| | Sex | 2.4 | 1.56 | 2.08 | 1 | 0.149 |
| | Side | -1.6 | 1.24 | 1.47 | 1 | 0.225 |
| | $SlopeSL = \beta_0 + \beta_{side} + \beta_{sex} + \epsilon$ | | | | | |
| **Slope SL** | Intercept | 3.7 | 0.58 | | | <0.001 |
| | Sex | 0.3 | 0.78 | 0.14 | 1 | 0.704 |
| | **Side** | **1.4** | **0.50** | **5.79** | **1** | **0.016** |

| | | | | | | |
|---|---|---|---|---|---|---|
| | $MeanWE = \beta_0 + \beta_{side} + \beta_{sex} + \epsilon$ | | | | | |
| **Mean WE** | Intercept | -1.9 | 0.04 | | | <0.001 |
| | Sex | 0.1 | 0.05 | 2.20 | 1 | 0.1383 |
| | Side | 0.0 | 0.03 | 0.19 | 1 | 0.6607 |

| | | | | | | |
|---|---|---|---|---|---|---|
| | $MeanME = \beta_0 + \beta_{side} + \beta_{sex} + \epsilon$ | | | | | |
| **Mean ME** | Intercept | -35.2 | 0.69 | | | <0.001 |
| | Sex | 1.3 | 0.79 | 2.45 | 1 | 0.1175 |
| | **Side** | **-1.9** | **0.79** | **4.90** | **1** | **0.0268** |

**Table S2. Linear mixed effects model outputs for the acoustic results over song development.**

| | Model parameter | Estimate of slope | Std. error | F value | DF | p |
|---|---|---|---|---|---|---|
| | $PTP_b = \beta_0 + \beta_{age}*age + \beta_{sex} + \beta_{side} + \epsilon$ | | | | | |
| **$PTP_b$** | Intercept | 0.812 | 0.090 | | | <0.001 |
| | Age | 0.001 | 0.001 | 1.307 | 1 | 0.257 |
| | Sex | -0.036 | 0.061 | 0.347 | 1 | 0.558 |
| | Side | 0.067 | 0.060 | 1.249 | 1 | 0.268 |
| | $PTP_t = \beta_0 + \beta_{age}*age + \beta_{sex} + \beta_{side} + \epsilon$ | | | | | |
| **$PTP_t$** | Intercept | 0.368 | 0.052 | | | <0.001 |
| | **Age** | **-0.002** | **0.001** | **5.287** | **1** | **0.025** |
| | **Sex** | **-0.095** | **0.035** | **7.212** | **1** | **0.009** |
| | Side | -0.049 | 0.035 | 1.987 | 1 | 0.164 |
| | $PTP_{icas} = \beta_0 + \beta_{age}*age + \beta_{sex} + \beta_{side} + \epsilon$ | | | | | |
| **$PTP_{icas}$** | Intercept | 0.147 | 0.066 | | | 0.031 |
| | **Age** | **0.003** | **0.001** | **11.083** | **1** | **0.001** |
| | Sex | 0.014 | 0.045 | 0.093 | 1 | 0.762 |
| | Side | 0.049 | 0.044 | 1.2E+04 | 1 | 0.274 |

| | Model parameter | Estimate of slope | Std. error | LL $\chi^2$ | DF | p |
|---|---|---|---|---|---|---|
| | $Minf_o = \beta_0 + \beta_{age}*age + \beta_{sex} + \beta_{side} + \epsilon$ | | | | | |
| **Min $f_o$** | Intercept | 482.1 | 30.3 | | | <0.001 |
| | Age | 0.26 | 0.4 | 0.4 | 1 | 0.510 |
| | Sex | -24 | 19.1 | 1.6 | 1 | 0.209 |
| | **Side** | **64.14** | **7.0** | **79.0** | **1** | **<0.001** |
| | $S1f_o = \beta_0 + \beta_{age}*age + \beta_{sex} + \beta_{side} + \epsilon$ | | | | | |
| **S1 $f_o$** | Intercept | 102.5 | 53.8 | | | 0.065 |
| | Age | 0.3 | 0.6 | 0.2 | 1 | 0.651 |
| | Sex | 18.3 | 37.9 | 0.2 | 1 | 0.630 |
| | Side | 29.6 | 26.6 | 1.2 | 1 | 0.270 |
| | $S2f_o = \beta_0 + \beta_{age}*age + \beta_{sex} + \beta_{side} + \epsilon$ | | | | | |
| **S2 $f_o$** | Intercept | 107.2 | 17.8 | | | <0.001 |
| | **Age** | **-0.5** | **0.2** | **4.9** | **1** | **0.027** |
| | Sex | -9.0 | 12.2 | 0.5 | 1 | 0.463 |
| | Side | -8.0 | 11.2 | 0.5 | 1 | 0.474 |
| | $MinSL = \beta_0 + \beta_{age}*age + \beta_{sex} + \beta_{side} + \epsilon$ | | | | | |
| **Min SL** | Intercept | 45.6 | 1.6 | | | <0.001 |
| | Age | 0.0 | 0.0 | 0.2 | 1 | 0.680 |
| | Sex | 1.1 | 1.1 | 0.8 | 1 | 0.374 |
| | Side | -0.8 | 0.7 | 1.2 | 1 | 0.271 |
| | $SlopeSL = \beta_0 + \beta_{age}*age + \beta_{sex} + \beta_{side} + \epsilon$ | | | | | |
| **Slope SL** | Intercept | 4.2 | 0.6 | | | <0.001 |
| | Age | 0.0 | 0.0 | 0.0 | 1 | 0.830 |
| | Sex | 0.1 | 0.4 | 0.0 | 1 | 0.874 |
| | **Side** | **0.6** | **0.3** | **4.3** | **1** | **0.037** |
| | $MeanWE = \beta_0 + \beta_{age}*age + \beta_{sex} + \beta_{side} + \epsilon$ | | | | | |
| **Mean WE** | Intercept | -1.748 | 0.045 | | | <0.001 |
| | Age | -0.001 | 0.001 | 3.3 | 1 | 0.070 |
| | Sex | 0.001 | 0.030 | 7E-04 | 1 | 0.979 |
| | Side | -0.026 | 0.015 | 3.0 | 1 | 0.084 |
| | $MeanME = \beta_0 + \beta_{age}*age + \beta_{sex} + \beta_{side} + \epsilon$ | | | | | |
| **Mean ME** | Intercept | -33.16 | 1.03 | | | <0.001 |
| | **Age** | **-0.03** | **0.01** | **3.98** | **1** | **0.046** |
| | Sex | 1.24 | 0.71 | 2.86 | 1 | 0.091 |
| | Side | -0.68 | 0.41 | 2.69 | 1 | 0.101 |

**Table S3. Linear mixed effects model outputs for MVM morphology.**

| | Model parameter | Estimate of slope | Std. error | F value | DF | p |
|---|---|---|---|---|---|---|
| | *MVC-LDC* = $\beta_0 + \beta_{age}$*age* $+ \beta_{sex} + \beta_{side} + \epsilon$ | | | | | |
| **MVC-LDC** | Intercept | 1106.5 | 40.8 | | | <0.001 |
| | Age | 0.4 | 0.5 | 0.6 | 1 | 0.446 |
| | Sex | 10.2 | 28.4 | 0.1 | 1 | 0.720 |
| | **Side** | **-85.7** | **24.9** | **10.0** | **1** | **0.002** |
| | *LDC-MDC* = $\beta_0 + \beta_{age}$*age* $+ \beta_{sex} + \beta_{side} + \epsilon$ | | | | | |
| **LDC-MDC** | Intercept | 641.8 | 62.5 | | | <0.001 |
| | Age | 1.0 | 0.8 | 1.7 | 1 | 0.191 |
| | Sex | -1.0 | 44.9 | 0.0 | 1 | 0.982 |
| | **Side** | **-61.9** | **24.1** | **5.9** | **1** | **0.015** |
| | *LDC Area* = $\beta_0 + \beta_{age}$*age* $+ \beta_{sex} + \beta_{side} + \epsilon$ | | | | | |
| **LDC Area** | Intercept | 24343.3 | 10734.9 | | | 0.030 |
| | **Age** | **274.2** | **134.1** | **3.9** | **1** | **0.048** |
| | Sex | -1740.8 | 7632.1 | 0.1 | 1 | 0.820 |
| | **Side** | **21356.0** | **5022.5** | **14.1** | **1** | **2.0E-04** |
| | *Bronchial Width* = $\beta_0 + \beta_{age}$*age* $+ \beta_{sex} + \beta_{side} + \epsilon$ | | | | | |
| **Bronchial Width** | Intercept | 1798.5 | 101.1 | | | <0.001 |
| | Age | 0.3 | 1.3 | 0.1 | 1 | 0.799 |
| | Sex | 100.8 | 72.5 | 1.9 | 1 | 0.172 |
| | Side | -59.4 | 39.3 | 2.2 | 1 | 0.139 |
